# Quantitative evaluation of growth rate and survival of planktonic yeast under dual-parameter photodynamic treatment

**DOI:** 10.1101/2025.11.09.687416

**Authors:** Nidia Maldonado-Carmona, Giacomo Insero, Franco Fusi, Giovanni Romano

**Affiliations:** Department of Biomedical Experimental and Clinical Sciences “Mario Serio”, Università degli Studi di Firenze, Firenze 50134, Italy

**Keywords:** photodynamic antimicrobial chemotherapy, Candida albicans, antifungal treatments, Rose Bengal, checkerboard model

## Abstract

Photodynamic Antimicrobial Chemotherapy (PACT) relies on the concomitant use of light and a photosensitizer molecule for microbial disinfection. Adding the dimension of light to the system increases the difficulty and variables to consider, yielding time and material-consuming experiments, with their reproducibility depending greatly on the disclosure of the light irradiation conditions. In the present work we analyse the effect of increasing irradiant exposures and rose bengal (RB) concentrations against *Candida albicans* ATCC 10231, in a checkerboard fashion. For this, we propose the use of the exponential phase coefficients, growth rate and initial number of cells, to assess the efficiency of the treatment. The coefficients are able to accurately describe the decrease of starting number of cells as consequence of the PACT effect, while also showing the metabolic burden of the presence of RB at high concentrations, with further hints that the incorrect combination of RB and radiant exposure decreases the growth rate without affecting the survivability. Our proposed methodology can then be used as a high-throughput screening method for new photosensitizers and new formulations, exploring their needs of radiant exposure for an efficient microbial disinfection.

## Introduction

The rise of microbial resistance towards our antimicrobial treatments is one of the biggest threats in modern medicine. Since the last decade, the World Health Organisation (WHO) and other policy makers have raised their concerns on the rise of antimicrobial resistance and their impact on the economy and the well-being of the world population. In 2023, the WHO released the global agenda for antimicrobial resistance in human health (World Health Organization 2024), where they disclose a list of 40 research and policy priorities to tackle antimicrobial resistance, focusing on prevention, diagnosis, treatment and care, and cross-cutting research. Additionally, the emergence of fungal infections and the rise of antifungal resistance has been addressed by the WHO fungal priority pathogens list (World Health Organization 2022), where they emphasize the need for research and development investments on innovative antifungal agents, focusing on strategies where there is no cross-resistance, and new chemical classes, new targets and new mode of action. The rise of fungal infections is burdened not only by the smaller variety of antifungal molecules, as compared with antibiotic molecules, but fungal infections will be exacerbated and will geographically expand, in the urgent context of the impending climatic crisis (Gadre *et al*. 2022). Thus, it is imperative for researchers to embrace the search for new antifungal and antimicrobial alternatives.

Photodynamic Antimicrobial ChemoTherapy (PACT) relies on the use of a photosensitizing molecule (PS), light irradiation and oxygen for the production of reactive oxygen species (ROS), mainly singlet oxygen (Type II reaction) or other ROS as superoxide anion and hydroxyl radicals (Type I reaction). The production of ROS has a deadly effect on the microorganisms exposed to it, resulting in an efficient therapy with hypothetical (Maisch 2015) low proneness for the development of resistance, and demonstrated lack of cross-resistance with drug-resistant strains (Kurosu *et al*. 2023). The reasons for PACT working as a “magic bullet” is addressed to a quick burst of ROS, which are radical species with the capacity to oxidize all the macromolecules, to which microorganisms are not ready to counteract. The stacking evidence of PACT efficiency is reflected in the number of clinical trials aiming for its exploitation, where from 203 current clinical trials related to photodynamic therapy, 72 (35%) are related to a disinfective application, but from these 41 (57% of PACT clinical trials) are related to dentistry applications(National Library of Medicine), which is understandable given the facility to irradiate the oral cavity. If PACT has not grown more as an alternative to antimicrobial treatments, it is due to the perceived difficulty to irradiate certain cavities, but mostly because of the heterogenous results obtained. Some reviews particularly address the heterogeneity of results obtained, when considering simple photosensitizers as methylene blue (Piksa *et al*. 2023) or rose Bengal (Fiegler-Rudol *et al*. 2025), with the panorama being more complex once considering the different light sources, microorganisms strains and treatments schemes used in the literature. Undeniably, the lack of harmonization is burdened by PACT being demanding in both time and consumables, when considering light and photosensitizers as two different dimensions.

Cellular death in a PACT scenario is a consequence of the chemotherapeutic effect and the photodynamic effect, with an ideal PS having low or null effect on cellular viability in the absence of light. Then, unlike a typical antibiotic, the photodynamic effect depends on the microorganisms receiving a lethal dose of ROS, which results from the conjunction of the proper light fluence rate and PS concentration. Traditionally, the decrease of viability as function of fluence rate and PS concentration is quantified using cell plating and colony forming unit (CFU) counting of the surviving microorganisms, a method that is time and resources consuming, while observing a high variability, both at interday and interlaboratory level (Beal *et al*. 2020). With the rise of efforts on making biomedical research more sustainable, several methods have arisen to try to alleviate and replace the burden of cell plating (Hazan *et al*. 2012; Meyer *et al*. 2023; Ebrahiminezhad *et al*. 2024), while relying on simple methodologies such as optical density, flow cytometry and micrographies analysis to quantify the number of alive microorganisms in sample.

Our proposed strategy consists of a comprehensive study of the effect of both light dose and PS, using planktonic *Candida albicans* as a model. We propose the analysis of the growth curves, and their exponential growth coefficients, as the parameters to define the efficiency of the treatment. Our proposed methodology was validated with samples starting with different starting cell densities, yielding a good prediction on the reduction of cells and supporting the robustness of the approach. Additionally, our proposed strategy brings into light the role that the growth rate could bring for researchers to further understand the effect of photodynamic therapy and their individual components. Furthermore, our strategy is useful for other physical disinfection methods, such as heating, pressure and gamma radiation.

## Materials and Methods

### Reagents, strains and culture media

Rose Bengal (RB) was acquired from Sigma Aldrich. *Candida albicans* ATCC 10231 was kindly provided by the Careggi Hospital, at the Università degli Studi di Firenze, and kept in a 30% glycerol stock at −80°C. All solutions were prepared in miliQ water. Glucose (VWR, 240 g L^−1^) was sterilized through filtration (0.22-micron filters, VWR). Sabouraud’s dextrose 4% agar (VWR, SA, peptone 5 g L^−1^, tryptone 5 g L^−1^, dextrose 40 g L^−1^, agar 15 g L^−1^) BHI2X-g media (VWR, brain heart infusion solids 35 g L^−1^, peptones 20 g L^−1^, glucose 44 g L^−1^, NaCl 10 g L^−1^, Na_2_HPO_4_ 5 g L^−1^) and BHI-g (VWR, brain heart infusion solids 17.5 g L^−1^, peptones 10 g L^−1^, glucose 22 g L^−1^, NaCl 5 g L^−1^, Na_2_HPO_4_ 2.5 g L^−1^), phosphate saline buffer (PanReac AppliChem, ready to use tablets pH 7.4, PBS, 1 tablet per 100 mL of buffer) were autoclaved before use. DMSO used for microbiological experiments was acquired from VWR and used without further treatment.

### Irradiation setup, light dose calculation and correction

For the PACT experiments, a 528 nm laser (Necsel Green NovaLum, VHP) source was coupled into a multimode 500 µm core fibre. The fibre output was collimated by using an off-axes parabolic silver mirror with 15 mm focal length, obtaining an optical beam with a waist of around 4 mm at approximately 10 cm from fibre end. The irradiation spot size is controlled by an adjustable iris positioned near the irradiation place, set to a 6 mm diameter. This configuration ensures that the optical beam fully fits within a single 96-well plate. After passing through the iris, the light spot shows sharp edges with negligible diffraction features. Laser power was measured with a calibrated power meter (Ophir Nova Display equipped with a PD300-UV-SH sensor head) and converted into irradiance by dividing it by the 6-mm diameter laser spot area. Experiments were performed using a radiant exposure ranging from 0.1 up to 50 J cm^−2^ obtained by multiplying the irradiance for the light delivery time, which ranges in between 7–160 s interval. Irradiation was conducted at room temperature, with ambient illumination of the order of 1 µW cm^−2^ which was considered negligible with respect to the laser irradiance.

The radiant exposure was corrected to better express the match between the used light source and the absorption capacity of the photosensitizer, as described by Schaberle (Schaberle 2018), allowing a better inter-experimental comparability when different light sources are employed. The light dose correction (LDC) factor for RB was calculated for RB dissolved in a mixture of DMSO 10% in PBS, a current mixture for non-soluble photosensitizers. The absorption spectra of RB 10 µM was measured, using a FL6500 (PerkinElmer) spectrophotometer in a 1 cm quartz cuvette. The emission spectrum of the 528 nm laser was acquired with an AvaSpec-ULS spectrophotometer (Avantes). The normalized absorption and emission spectra were matched along their wavenumber (Supplementary Figure 1), resulting in an LDC of around 0.47 for RB in DMSO 10%.

### Photodynamic activity

Upon utilization, a smear of *C. albicans* frozen stock was spread on Sabouraud’s agar and incubated at 30°C for 24 hours. The obtained plate was stored in the fridge and used within five days. On the experimental day, a smear of 10 single colonies was suspended in 1 mL of PBS and incubated one hour at 37°C, with constant stirring (50 rpm). Cells were then diluted into 15 mL of PBS to an OD_600_ of 0.3, which roughly corresponds to 2 x 10^6^ CFU/mL. Typically, RB were prepared in a stock solution of 1 mM in DMSO, which was kept in the dark at −20 °C. Then, the stock solution was diluted in DMSO to 10X the intended final concentrations. This working solution was then diluted to 2X the intended concentration, in a mixture 20:80 DMSO:PBS, from which 60 µL were placed in a 96-well plate (VWR), where it was mixed with 60 µL of the yeasts suspension, resulting in a 1X RB 10:90 DMSO:PBS suspension. Controls included cells exposed to i) only RB at different concentrations, ii) only light irradiation at different irradiance exposures, and iii) without either RB or light exposure; as blanks, samples with RB but without yeasts were prepared and the blanks were not irradiated. Controls without RB were prepared yielding a 10:90 DMSO:PBS mixture. The cells were exposed to RB (37 °C, 50 rpm) for 45 minutes, and were immediately exposed to the irradiation process. After the irradiation process, 20 µL of each well were taken and spot-plated (5 µL per spot) in Sabouraud agar plates. Then, 100 µL of BHI2X-g media were added to all wells yielding a mixture of BHI with glucose 20 g L^−1^and 5% DMSO. The plate was sealed with parafilm and brought to the Tecan multi-well plate reader. *C. albicans* growth was followed at 600 nm, measuring the plate every 10 minutes (37 °C, linear shaker, 5 seconds, 1 mm amplitude), up to 24 hours. Spotted Sabouraud agar plates were incubated at 37 °C overnight, followed by CFU counting.

### Validation of the cell reduction assessment method: preparation of cultures

Three yeast suspensions were prepared at a starting OD_600_ of 0.3, 0.6 and 0.9, as measured in a common OD reader (Implen). Then, the suspensions were serially diluted in PBS, decreasing the cell densities from 10^−1^ to 10^−6^. The different cell suspensions were dispensed (100 µL) in a 96-well plate by triplicate and fed with 100 µL of BHI2X-g. The plate was sealed with parafilm and brought to the Tecan multi-well plate reader. *C. albicans* growth was followed at 600 nm, measuring the optical density of each well every 10 minutes (37 °C, linear shaker, 5 seconds, 1 mm amplitude), up to 24 hours. In addition, the yeasts suspensions were serially diluted in PBS, plating 100 µL in Sabouraud agar plates and spreading the suspension with a T-spreader, and incubated for 16 hours at 37 °C for CFU counting.

### Cell reduction and growth rate assessment

Data processing was done using MATLAB 2025a. An example of the scripts used can be found at Supplementary Material 1.

Raw optical density 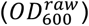, was obtained from the multiplate reader as an .csv file, with acquisitions both in time (rows) and space (columns). An example of the process can be found in Supplementary Figure 2.

To account for RB absorption at the monitoring wavelength (600 nm), adequate controls were prepared with decreasing concentrations of RB and monitored across time, to reduce any systematic errors due to prolonged measurements. Upon completion of the growth curve reading, samples were grouped by RB concentration and the optical density (*OD*) of their respective blanks were subtracted.

To extract the growth descriptors, the growth rate (*μ*) and the calculated starting optical density 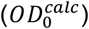, we calculated numerically the time derivative of *OD*(*t*) (*dOD*/*dt*) as *δOD*_600_/*δt* with *δt* = 0.167 hours (10 minutes) (see Supplementary Figure 2A and 2B). This was used to assess the presence of a local maxima, indicating an inflection point that represents the transition between the exponential and stationary growing phase. Here, we consider the exponential phase where the time-derivative values range from 20% to 80% of the found local maxima, allowing us to neglect the transition between lag-exponential and exponential-stationary phases. The chosen values to consider the exponential phase (20-80%) were arbitrarily chosen, but our observations demonstrate that as long as the initial lag-phase is not considered, the method yields a robust prediction (Supplementary Figure 3), thus other values could be used, depending on the analysed sample. The OD data corresponding to the exponential phase only were reported in a semilogarithmic plot and fitted with the linear function:

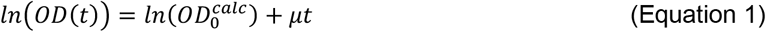

to give the growth rate *μ* and *OD*^*calc*^ for each sample (Supplementary Figure 2C). In the case of the photodynamic effect, we can define the survival rate as:

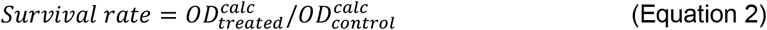

As a point of comparison, we include the SGT methodology, which was previously found to be successful to predict reduction of bacterial and fungal planktonic cells (Hazan *et al*. 2012). The ΔSGT value was calculated following the equation

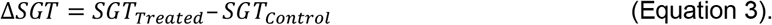

## Results and discussion

### Exponential growth parameters as predictors for viability in *Candida albicans*

Under ideal conditions, the growth of both planktonic yeast and bacteria is exponential, given that the cells duplicate themselves at a characteristic time. Planktonic growth passes through three distinctive phases: i) lag phase, where cells adapt to new conditions and prepare for duplication, ii) exponential phase, where cells duplicate and are metabolically active, and iii) stationary phase, where lack of nutrients and waste accumulation pushes to a slow down on the growth rate (Figure 1A). The exponential phase can be described by the classical equation

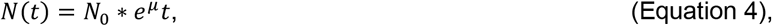

where *N*(*t*) is the number of cells as function of time *t, N*_0_ is the starting number of cells, and *μ* is a characteristic growth rate, which is specific for the given conditions (i.e. temperature, culture media, etc.). In terms of optical density, it is acknowledged that there is a proportionality between *OD*(*t*) and *N*(*t*), during the exponential phase.

**Figure 1.**
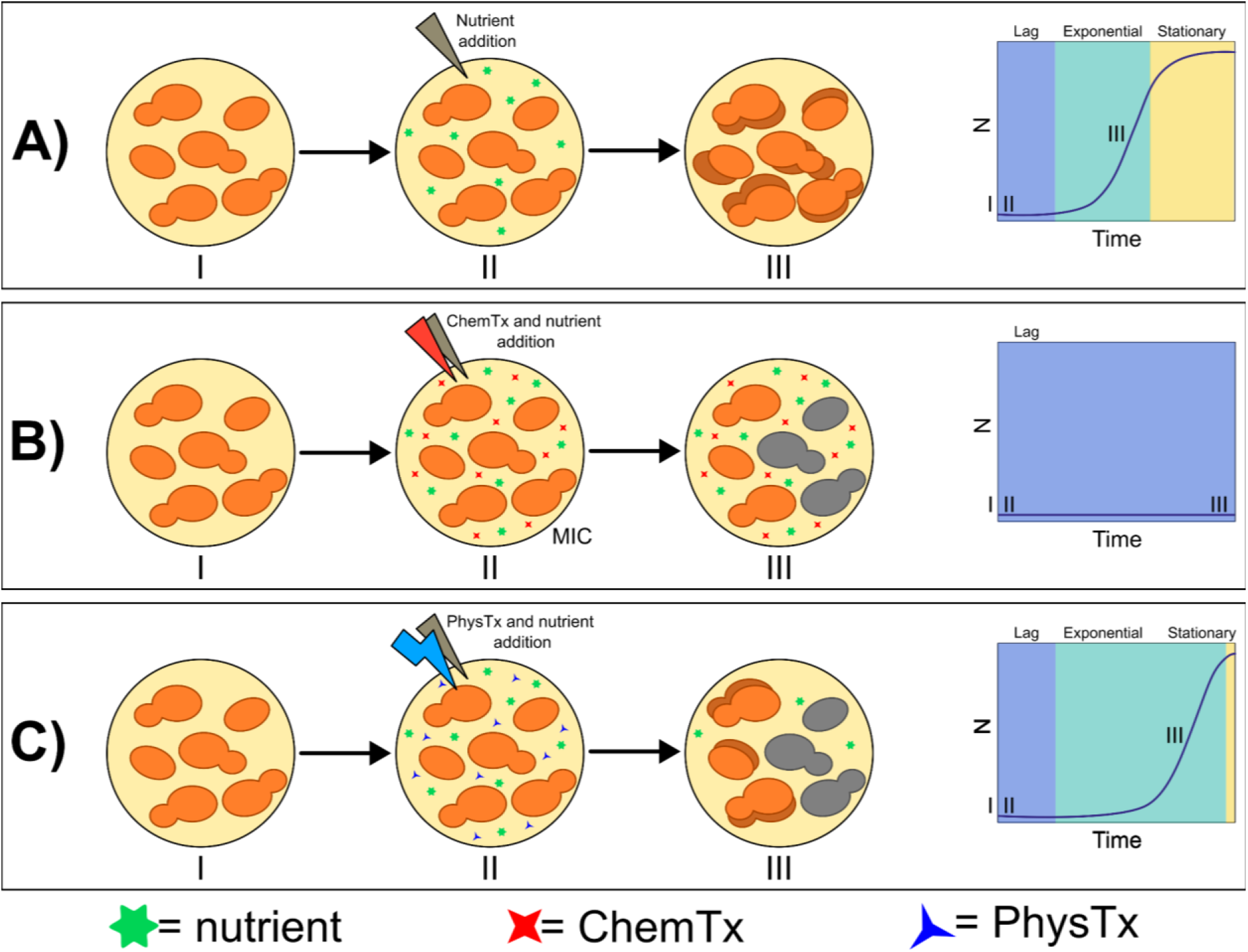
Microorganisms’ growth under **(A)** ideal conditions, **(B)** Chemotherapeutic treatment (ChemTx) or **(C)** Physical treatment (PhysTx). Under ideal conditions **(A)**, an initial number of microorganisms (i) are able to replicate when their nutrimental needs are met (ii), with their populations following a typical lag or adaptive phase, followed by an exponential phase (iii) and a stationary phase, where waste accumulation and the lack of nutrient stalls the growth. When microorganisms are exposed to an inhibitory concentration of ChemTx **(B)**, growth is stalled due to a combination of microstatic and microcidal events, as long as the MIC is kept. When microorganisms are exposed to a PhysTx event **(C)**, a number of cells die as result of the treatment at a given time (ii), with the surviving cells being able to recover and eventually replicate, delaying the onset of the (iii) exponential phase.

In the simplified case where a sample with a given starting experimental optical density 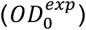 is composed of only alive cells, we can assume that 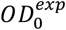 is proportional to *N*_0_ and we can state that the exponential phase of this population will follow Equation 1, where 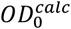 is found as the intercept of the linear fit of *ln*(*OD*(*t*)) and which is proportional to the starting number of cells (*N*_0_) and matches the experimental value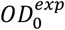.

Physical disinfection methods are widely used for the disinfection of food, medical equipment, and medical graftings (Yan *et al*. 2021), using technologies as diverse as ultra-high pressure (D’Incecco *et al*. 2018; Nikparvar *et al*. 2021), heating (Silva *et al*. 2023), gamma radiation (Annamalai *et al*. 2020), ultraviolet radiation (Miller, Linnes and Luongo 2013) and PACT (Maldonado-Carmona *et al*. 2020). Chemotherapeutic approaches (Figure 1B) affect the microorganisms through their constant presence at a given effective concentration (minimal inhibitory concentration, MIC) where microorganisms growth is inhibited, due to a stall of the microorganism replication or due to cellular death. Physical treatments are differentiated from chemotherapeutic approaches as the treatment is applied over a well-defined and finite time interval (Figure 1C), and not as a constant lethal exposure across time. Once the physical treatment is over, microorganisms are not exposed to deadly conditions. Then, samples exposed to a physical treatment are a mixture of alive and dead microorganisms, without further death happening, with the surviving microorganisms being able to replicate if their nutritional conditions are met. In this case, 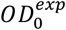 is no longer proportional to *N*_0_, as 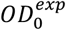 is a mixture of both dead and alive cells. Using our previous assumptions, we can model the post-event cell growth by invoking the same exponential model, considering that there is a decrease in *N*_0_, compared to an unaffected sample. Assuming that the growth rate of the surviving cells has been unaffected by the physical event, the exponential phase onset will be delayed, with the intercept of *ln*(*OD*(*t*)) indicating a lower 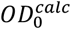, which would reflect the reduction of *N*_0_ after the physical treatment.

### Growth rate and prediction of initial number of cells in *Candida albicans*

Then, after a physical treatment, we can consider that any changes in *OD*(*t*) are due only to the presence of surviving cells. To assess these assumptions and validate our proposed method we artificially decreased the concentration of cells, through serial geometrical dilutions (Figure 2A). As the growth curves are the result of both the growth rate and the initial number of cells, decreasing the starting number of cells has an immediate effect on the growth kinetics, showing a delay on the growth onset as a function of time. In the simplified case where the starting number of cells, expressed in optical density 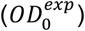, has been artificially decreased and the growth rate is *a priori* the same for all the conditions, we observe that the onset of the exponential phase is delayed. Applying our proposed methodology, we were able to extract the exponential growth parameters, which accurately fit the exponential phase, although the confidence of the fit decreases for samples with the lowest 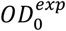 values. The extracted parameters were then compared with the ground truth, the actual number of cells present at the beginning of the experiment.

**Figure 2.**
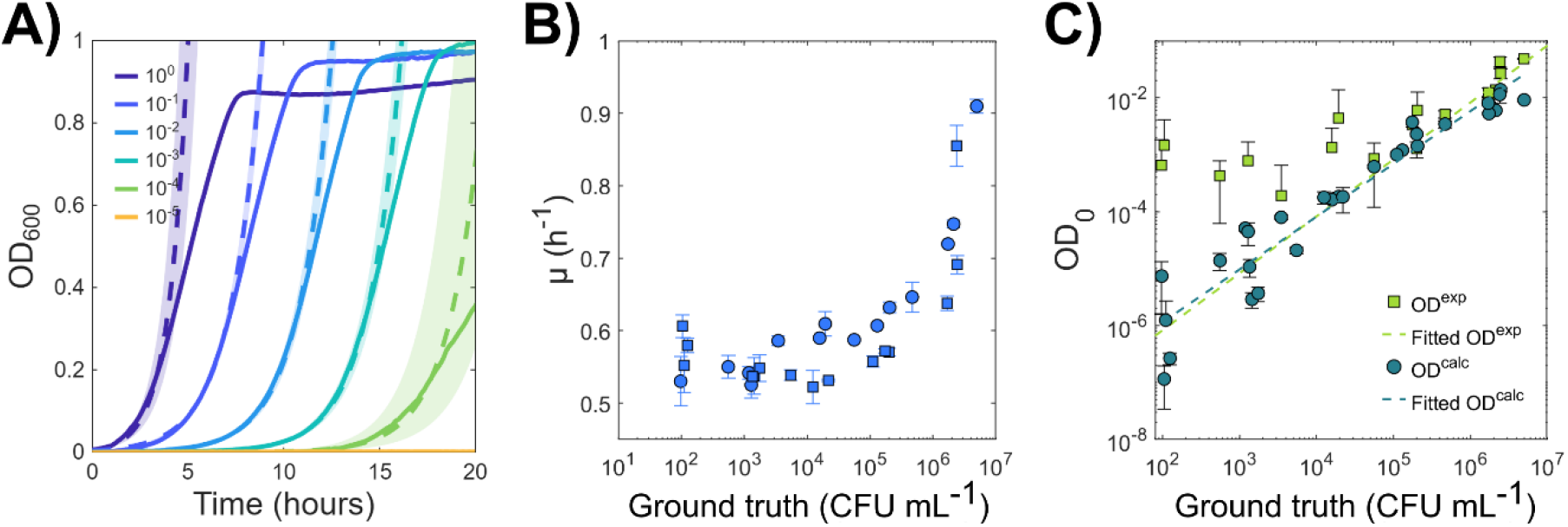
Data extracted from the growth curves. **A)** Matching of the observed growth curves (solid lines) and their estimation (dashed lines) obtained from the equation 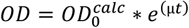. Shadowed areas depict their respective confidence bound at 95%. **B)** Growth rates calculated for different cell suspensions with different starting cell densities. Solid symbols and error bars depict the mean and standard deviation of three technical replicates, while the squares and circles represent the results of two independent experiments. **C)** *OD*_0_ as a function of decreasing cell densities. Solid symbols and error bars depict the mean and standard deviation of three technical replicates. Light green squares depict the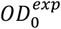, as determined by the multiwell plate reader, with the light green dashed line fitting 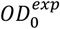 while assuming a slope = 1, which would correspond to a perfect geometrical dilution. Dark green circles depict the 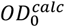, as extracted from the growth curves, with the dark green dashed line fitting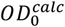.

When decreasing the starting cell density, we explored the hypothesis that the growth rate may not be constant across different cell densities, especially when considering that *C. albicans* is sensitive to quorum sensing (Hornby *et al*. 2001; Kruppa 2009), although the reports in the literature have focused more on *C. albicans* morphological shifting upon changes in the starting cell densities, with barely no comments towards changes in the growth rates. When we analysed the growth rate as a function of decreasing cell densities, we observed an increase in the growth rate for samples above 5 x 10^5^ CFU/mL, rising from around 0.6 up to 0.9 h^−1^ at the highest cell densities (Figure 2B). Previously (Hornby *et al*. 2001), it has been reported that *C. albicans* cultures starting with high cell densities (10^7^ CFU/mL) have a higher prevalence of the budding yeast morphological state, when compared culture grown under the same conditions but with lower starting cell densities (10^5^ CFU/mL). Although researchers did not purposely analyse the growth rate of their resulting curves, it was noted a change on the shift from exponential to stationary phase, at around 12 and 48 h respectively, while showing a higher growth rate for high density cell cultures, simply observed as a difference in the step of the slope. Our results are compatible with the hypothesis that starting with cell densities above 5 x 10^5^ CFU/mL may provoke the accumulation of quorum sensing molecules (QSM), which are likely to trigger morphological and behaviour changes, while cultures starting with lower cell densities (< 5 x 10^5^ CFU/mL) show a steady growth rate at the used culture conditions.

On the other hand, we observed a linear relation between the 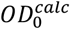 and the ground truth (Figure 2C) along the range of 10^2^-10^7^ CFU/mL, although below 10^3^ CFU/mL we could observe a decrease in the prediction capacity, as observed by a broadening of the confidence bounds. Below 10^2^ CFU/mL, growth was not found for all the triplicates and the number of cells counted in the spot method were found to be below 20 cells/plate, which is considered the lower detection limit of the CFU counting method. Remarkably, although the growth rate was found to variate across different starting optical densities, especially at cell densities above 5 x 10^5^ CFU/mL, the 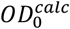 follows the expected behaviour of 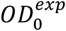, as determined through the few available samples where the experimental starting optical density is detected by the multi-well plate reader (Figure 2C, light green dashed line).

### Rose Bengal photodynamic effect against *Candida albicans*

Rose Bengal (RB) is a well described photosensitizer with described photoactivity against bacteria (Durkee *et al*. 2020; Gavara *et al*. 2021; Hirose *et al*. 2021; Kurosu *et al*. 2023; Trevizani Rocchetti *et al*. 2023), viruses (Cossu *et al*. 2016; Kingsley *et al*. 2018; Eshaghi Gorji and Li 2022) and fungi (Gavara *et al*. 2021; Bekmukhametova *et al*. 2024; Aydinoglu *et al*. 2025), with ranging applications in dentistry, agriculture and medicine. RB is an old (Ito 1978) and well-known molecule, approved by FDA as a colouring agent and with described innocuity in the dark. Together with its high solubility in aqueous media and high singlet oxygen quantum yield (Φ_(1O2, H2O)_ = 0.75) (Murasecco-Suardi *et al*. 1987), it is easily described as a “perfect” photosensitizer, resulting in an ideal molecule to test our method.

Given the wide use of RB across time, it is also an example of the wide variability of results found in PACT for a single molecule^8^ (Table 1). The wide range of described efficiencies is not really surprising, as PACT relies on chemical (i.e. compound, solubility, pK_a_, etc.) and physical (i.e. irradiance, power, wavelength of irradiation, light source used, full width at half maximum (FWHM) of the used light source, irradiation time, etc.) parameters for providing an effect on a biological system, which response to the treatment will also depend on the starting conditions of the subject (i.e. planktonic state, biofilm lifestyle, culture media, strain, etc.).

**Table 1.** Results found in the literature, where RB has been used as a photosensitizing agent against planktonic cultures of *Candida albicans*.

| Ref. | Culture conditions |  |  | RB |  | Irradiation conditions |  |  |  |  |  | Results |  |
| --- | --- | --- | --- | --- | --- | --- | --- | --- | --- | --- | --- | --- | --- |
|  | Strain | Inoculation media | Cell density | Concentration | Delivery method | Light source | Irradiation time point | Wavelength of irradiation | Irradiance | Irradiation time | Radiant exposure | Methodology | Viability decrease |
| (Costa et al. 2012) | *C. albicans* ATCC 18804 | PBS | OD 0.284 ($10^6$ CFU/mL) | 10 μM | PBS | LED | After 5 minutes of incubation | 455 +/- 20 nm | 526 mW cm<sup>-2</sup> | 180 s | 95 J cm<sup>-2</sup> | Counting colony-formin units | 1.58 +/- 0.35 log reduction<sup>15</sup> |
| (Freire et al. 2014) | *C. albicans* ATCC 18804 | PBS | OD 0.284 ($10^6$ CFU/mL) | 6.25 μM | PBS | LED | After 5 minutes of incubation | 532 +/- 10 nm | 237 mW cm<sup>-2</sup> | 180 s | 42.63 J cm<sup>-2</sup> | Counting colony-formin units | ~4.5 log reduction |
| (Wen et al. 2017) | *C. albicans* CEC 749 | PBS | OD 0.6 ($10^7$ CFU/mL) | 10 μM | PBS | White lamp with a band-pass filter | After 30 minutes of incubation | 540 +/- 158 nm | 100 mW cm<sup>-2</sup> | ND | 20 J cm<sup>-2</sup> | Counting colony-formin units | 0 log reduction |
| (Valkov, Zinigrad and Nisnevitch 2021) | *C. albicans* ATCC 90028 | Saline solution 0.9% | OD 0.1 ($10^6$ CFU/mL) | 50 μM | Saline solution | White luminescent lamp | ND | 400 – 700 nm | 1.9 mW cm<sup>-2</sup> | 15 minutes | 1.7 J cm<sup>-2</sup> | Counting colony-formin units | ~ 6 log reduction |

One of the most frequently found difficulties is how researchers measure and express light irradiation. Many times, researchers will not include all the relevant information needed to reproduce or properly compare the irradiation conditions used, which results particularly complicated when using broad light sources instead of laser lines. This is highlighted in Table 1, where the efficiency of the treatment ranges from 0 to six orders of magnitude, although it cannot be discarded that the differences are due to biological differences between the tested strains. Furthermore, it has been suggested (Schaberle 2018) that the emitted fluence rate needs to be corrected with a degree of overlap between the emission spectra of the light source and the absorption spectra of the photosensitizer, as most of the times there is not a perfect match between the light source and the photosensitizer absorption peaks (Supplementary Figure 1).

The difficulty to test all the spectra of variables that can influence an effective photosensitization is burdened by the lack of screening methods for photosensitizing agents, methods that should consider adding light as a second dimension. In this sense, we can consider RB concentration and radiant exposure as two interactive variables and we can assess their synergic effect using a checkerboard model (Figure 3A). The checkerboard model is widely used in microbiology and pharmaceutical sciences to determine if two molecules have a synergistic effect against axenic cultures (Cokol-Cakmak and Cokol 2019), with this approach determining a synergic MIC. We propose using the checkerboard model, where photosensitizer concentration (0-50 µM) and radiant exposure (0-50 J cm^−2^) are treated as the two interacting variables. The irradiation conditions account for an irradiance between 0 and 0.3 W/cm^2^ (6 mm diameter spot), allowing the delivery of up to 50 J cm^−2^in less than three minutes, with the irradiation time being negligible compared to the typical yeast doubling time (∼1.2 hours). The irradiation time was measured for each individual well and the light dose correction for RB was applied (Supplementary Figure 1). Furthermore, the irradiation step was performed in the absence of growth media, in order to avoid the presence of replicating cells and non-specific absorber molecules during the irradiation process.

**Figure 3.**
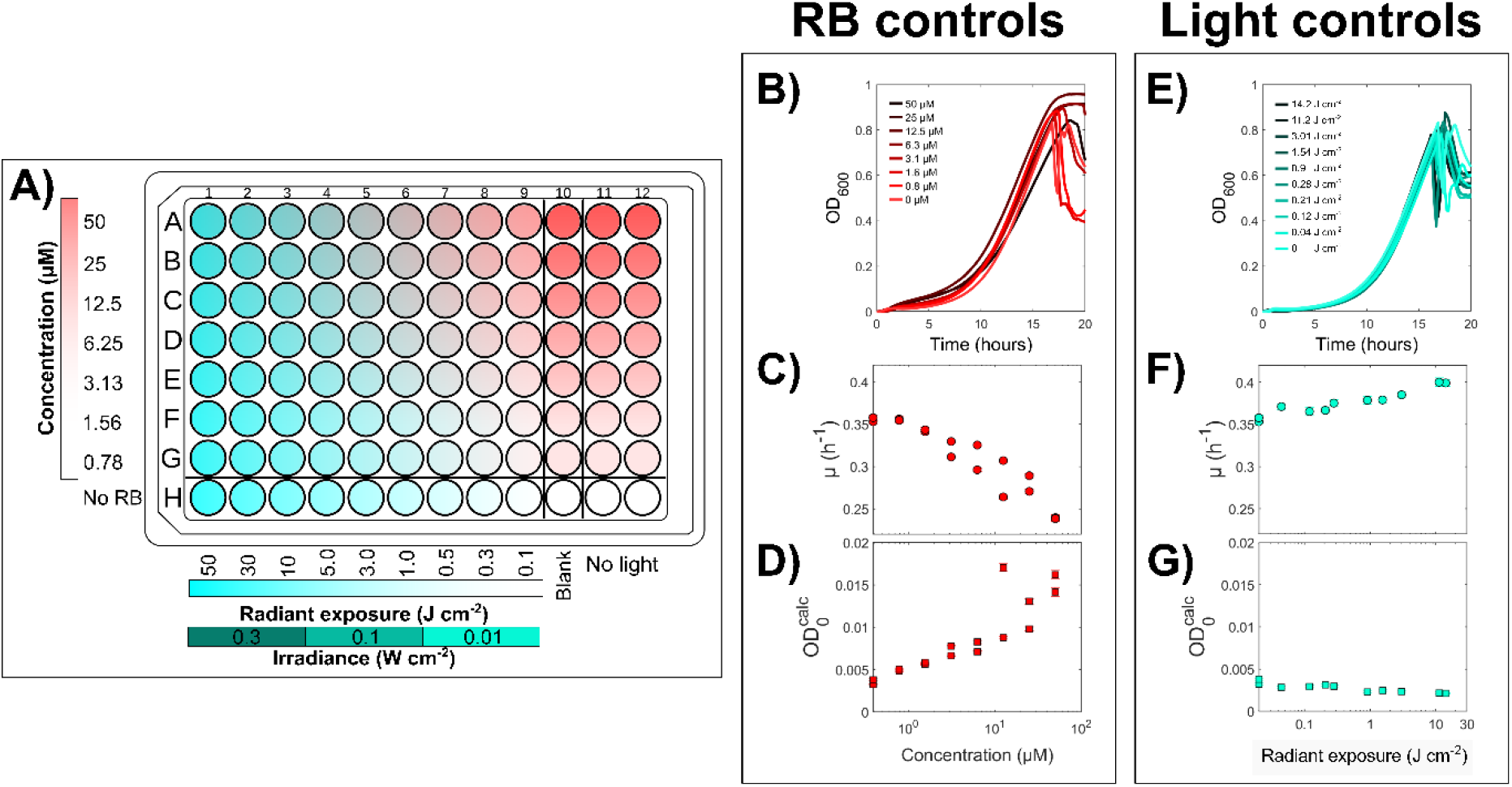
Experimental design and growth curves found in the checkerboard model. **A)** Checkerboard design with increasing RB concentrations and increasing radiant exposures, as consequence of increasing irradiances and irradiation times. **B)** Growth curves of controls exposed to increasing concentrations of RB, including the non-RB exposed control. **C)** Growth rate and **D)** 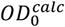 of controls with increasing concentrations of RB. Solid symbols represent the predicted value, with the confidence of fit at 95% depicted as whiskers. **E)** Growth curves of controls exposed to increasing radiant exposures, including the non-light exposed control. **C)** Growth rate and **D)** 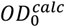 of controls with increasing radiant exposures. Solid symbols represent the predicted value, with the confidence of fit at 95% depicted as whiskers.

The growth curves of the two groups of controls, RB controls (Figure 3B-D) and light controls (Figure 3E-G) resemble the shape of the growth curves of samples not exposed to either light or RB. Both groups show affectations at the stationary growth level. Upon extraction of the growth parameters, it is brought into evidence that separately light irradiation has no effect on the development of the exponential phase (Figure 3E-G), while the presence of RB seems to decrease the growth rate and increase the 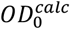. Although it is technically possible that some wells have a higher number of cells than the non-treated controls, this would be due to experimental variability, which is not what we observe, where the 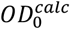 at the highest doses of RB is three times higher with respect to the control, and then it could be considered an artifact of our method. However, such behaviour was not found during the validation process, possibly being exclusively due to the presence of RB. On the other hand, it has been previously described that high concentrations of RB have a mild cytotoxic effect in eukaryotic cells (Lee *et al*. 1996; Mousavi *et al*. 2009). In our case, we observe that the presence of RB acts as a mild cytostatic, decreasing the growth rate with a dose-response behaviour as the growth rate at 50 µM decreased by 30 % with respect to the non-RB controls (Figure 3C).

Remarkably, the exposure to light only seems to have an effect on the growth rate, as cells seem to have increased their growth rate, with respect to the dark controls. Certain reports indicate that oil producer yeasts such as *Rhodotorula glutinis* (Yen and Zhang 2011) and *Rhodosporidium toruloides* (Pham *et al*. 2020) increase its growth rate as a function of light exposition, which has potential uses for lipid and carotene production. However, both reports were done under different irradiation conditions, and using broad wavelength light sources. This could possibly imply that *C. albicans* is capable of regulating its metabolic functions in the presence of light, although the photomodulation and their mechanisms of *C. albicans* has not been explored, to the knowledge of the authors.

The joint exposition of the cells to RB and light leads to a shift on the onset of the exponential phase (Figure 4A), with the shift responding to the dose of RB and radiant exposure, as observed when artificially decreasing the number of cells. When the optical density after 24 hours was no different than the blank, the 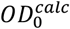 was assumed to be the lower limit of detection of our method (*lld* = 1.11 x 10^−6^), and the growth rate was not calculated. The different growth parameters were calculated, including 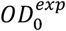 (Figure 4B), final OD (Figure 4C), the growth rate (Figure 4D) and 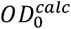 (Figure 4E).

**Figure 4.**
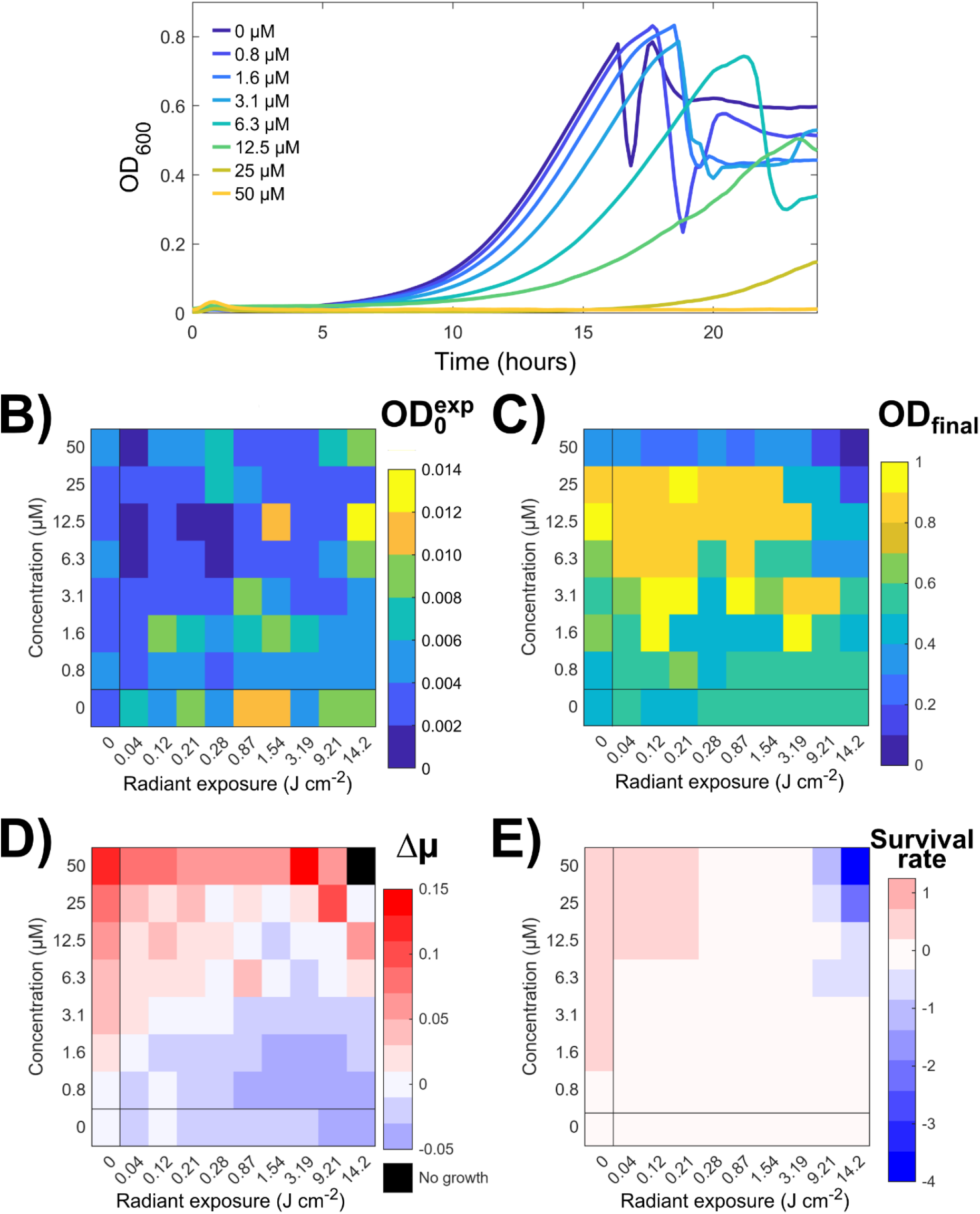
Photodynamic effect on *C. albicans*. **A)** Growth curves of C. albicans exposed to 14.2 J cm^−2^ and different concentrations of RB. **B)** Starting experimental optical density 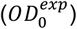, as detected by the multiwell plate reader. **C)** Optical density after 24 hours of growth (*OD*_*final*_), as detected by the multiwell plate reader. **D)** Growth rate change (Δμ, calculated as Δμ = μ_*control*_ − μ_*sample*_). **E)** Survival rate, calculated as 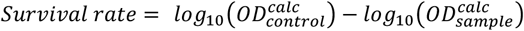. Samples where no growth was detected were assigned with the lowest value that our method is able to determine, the lower limit of detection (*lld* = 1.11 x 10^−6^), as estimated from Figure 2C.

The observed starting optical density, 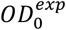, was compared across the different conditions, without finding evidence that 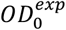 changes as a consequence of the different photodynamic conditions (Figure 4B). This is not surprising, as even if cells were killed, the sensitivity of the multiwell plate reader would not permit us to determine whether the number of cells has been reduced. On another hand, we were able to observe that the OD at the end of the kinetics (t = 24 h, *OD*^*final*^) shows interesting patterns and defined sections in the checkerboard (Figure 4C), but lacking a dose-response logic. The explanation found is that for certain wells, the optical density decreases during the stationary phase, which could be due to accumulation of biomass on the edges of the wells. In any case, the stationary phase is not suitable for measuring the efficiency of the photodynamic treatment.

The combination of light and RB yields a wide range of effects, with our growth descriptors indicating that both growth rate (Figure 4D) and viability (Figure 4E) is being affected by the photodynamic therapy. As expected, the most radical effect is found at the highest RB concentrations and radiant exposures applied, where a lack of growth was found. The difference of the growth rate with respect to the controls not exposed to light nor RB, Δμ = μ^*control*^ − μ^*sample*^, allows us to identify two regions, where the growth rate is dominated by the presence of RB, marked by a decrease of the growth rate, or by the presence of light, marked by a slight increase on the growth rate. This pattern is broken in the regions where the cells were exposed to RB concentrations above 12.5 µM and radiant exposures above 3.19 J cm^−2^, where the growth rate decreases even further, with the biggest effect found at 50 µM and 3.19 J cm^−2^. In these conditions, we could hypothesize that the cells are being affected not only by the constant presence of RB, but subdue to affectations related to the photodynamic treatment. This suggests that under the given conditions, cells are damaged and are able to react accordingly. Upon infection in a human host, *C. albicans* is internalized by macrophages, which are then contained inside lysosomes, where superoxide anion and nitric oxide radicals are produced by NADPH oxidase and nitric oxide synthase, respectively (Arribas, Gil and Molero 2025). The response of *C. albicans* to the internalization includes i) the production of catalase, ii) a metabolomic reprogramming to a catabolic state (gluconeogenesis, fatty acid degradation), iii) expression of DNA damage repair and oxidative stress genes, iv) switch to hyphal growth, allowing escape from the macrophage and the restart of glycolytic growth (Lorenz, Bender and Fink 2004). It is plausible that, in a pathological scenario, the presence of ROS in the surrounding volume of *C. albicans* will affect the growth rate, especially as the transcriptomic data (Lorenz, Bender and Fink 2004) already suggests that the cellular cycle is stalled in the G1 phase.

Interestingly, although we are able to observe an intense effect on the growth rate upon the photodynamic treatment and their individual elements, this is not reflected in the changes on the survival rate. We found that only at the extreme conditions (i.e. the highest RB concentrations and the highest radiant exposures) there is a substantial decrease of viability above three orders of magnitude. When decreasing either the RB concentration or the radiant exposure, the efficiency of the treatment (Figure 4E) quickly decreases, with the immediate neighbouring wells yielding only one order of magnitude reduction, which was corroborated through spot-platting (Supplementary Figure 4). The fact that under non-lethal conditions the growth rate is affected but not the viability of the cells is something that should not be ignored: microorganisms’ survival may lead to tolerance and resistance. One of the best properties of PACT is its apparent lack of resistance, although in the latest years this has been brought into deeper discussion, with results showing that antimicrobial Blue Light, a specific type of PACT relying on endogenous photosensitizers and blue light irradiation, is able to create tolerance in Gram-negative bacteria (Rapacka-Zdonczyk *et al*. 2021). Although further studies are still needed, it is widely accepted that cells with low growth rate are less susceptible to antimicrobials (Pontes and Groisman 2019), and thus surviving cells with a diminished growing rate may be less susceptible for further treatment, in a clinical scenario. Thus, although an excellent photodynamic effect is found when using RB at 50 µM, cells that do not receive the lethal light dose not only will survive the photodynamic treatment, but also they may be less susceptible to further chemotherapeutic treatments, and could be even potentially primed for further PACT treatments. This is relevant for the exploitation of PACT in a clinical scenario, where cells may be found on three-dimensional arrangements (i.e. biofilms) that may prevent or diffuse the light irradiation, which could lead to resistance to antimicrobial drugs or prevent the absorption of other photosensitizer molecules.

The use of the growth rate as a descriptor, complements the information obtained through the survival rate of cells. Both descriptors are easily obtained with our proposed methodology, which otherwise would need two separated experiments to obtain. Then our methodology exploits information that is easily obtained by researchers, using tools and material that is normally found in a biology laboratory, while reducing the load of plastic usage. The burden of plastic consumption in microbiology, especially in assays that involve dilution and plating of multiple samples, has been brought into consideration by other researchers (Hazan *et al*. 2012; Meyer *et al*. 2023) with the current work helping in the look for protocols with better time, data and material resources consumption.

### Comparison with other methods

Our proposed method tries to circumvent the tedious process of counting colonies to determine the microorganism’s survival after a given treatment. Colony forming counting has been demonstrated to be a method with low reproducibility in interday and interlaboratory setups (Beal *et al*. 2020), while being time and material demanding. Alternative methodologies take in account the preparation time, the material consumption, the analysis time and the availability of material in a standard microbiology culture room. Some of the simplest analyses rely on optical density analysis, given the wide spread of multi-well plate readers in microbiology laboratories. A classical method is the one proposed by EUCAST (European Committee for Antimicrobial Susceptibility Testing (EUCAST) of the European Society of Clinical Microbiology and Infectious Diseases (ESCMID) 2003), which has been successfully adapted for the determination of the minimal inhibitory concentration of photoactive compounds of natural origin (PhotoMIC) (Fiala *et al*. 2021). The EUCAST method is designed for the evaluation of antibiotics and antifungals, as the microorganisms are constantly exposed to different concentrations of a chemotherapeutic drug. It proposes an end-point measurement, at the end of an incubation at 24 hours, with the concentration where growth is not found being considered as the minimal inhibitory concentration (MIC), assuming that the lack of growth can be due to i) a microstatic effect, where growth has been constantly inhibited but microorganisms are alive, or ii) a microbicidal effect, where the microorganisms have been killed. To differentiate, further testing is needed to determine the minimal microbicide concentration (MBC), usually needing higher concentrations to achieve a microbicidal effect. Results obtained from this test are considered semiquantitative as it is normally not reported the percent of inhibition found, but the concentration at which no growth was found. In this sense, only one condition found at Figure 4C would be considered as positive for growth inhibition, corresponding to 50 µM and 14.2 J cm^−2^, without further considerations on the biological effect on other wells.

Another method relying on optical density sets an arbitrary threshold of optical density and determines the time when this threshold is crossed, defined as the Starting Growth Time (SGT), which can be defined as the time when growth happens, with previous results showing a good sensitivity to predict the reduction in the number of bacteria present in a sample (Hazan *et al*. 2012). However, this method results useful while assuming that the growth rate is constant for all the samples and remains unaffected by a possible treatment, which may be the case for a healthy population artificially diluted (Figure 5A) but which is unlikely when testing an antimicrobial treatment. This was studied through the simulation of exponential curves with different starting number of cells (*N*_0_) and assuming different growth rates (Supplementary Material 3). When calculating the SGT for curves with different growth rates (Figure 5B), we can observe a change in the slope, limiting the predictive potential of this methodology to populations where all the conditions have the same growth rate. We extracted the SGT values from the data obtained for the PDT treatment of *C. albicans* with RB (Figure 5C), and compared them with the controls (Δ*SGT* = *SGT*_*treated*_ − *SGT*_*control*_). The results obtained reflect those found when analysing the 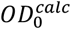 (Figure 4E), showing that affectations are not limited to the conditions where growth is absent. However, unless compared with a standard curve and assuming that the growth rate is constant along the different samples, a quantitative determination may be uncertain.

**Figure 5.**
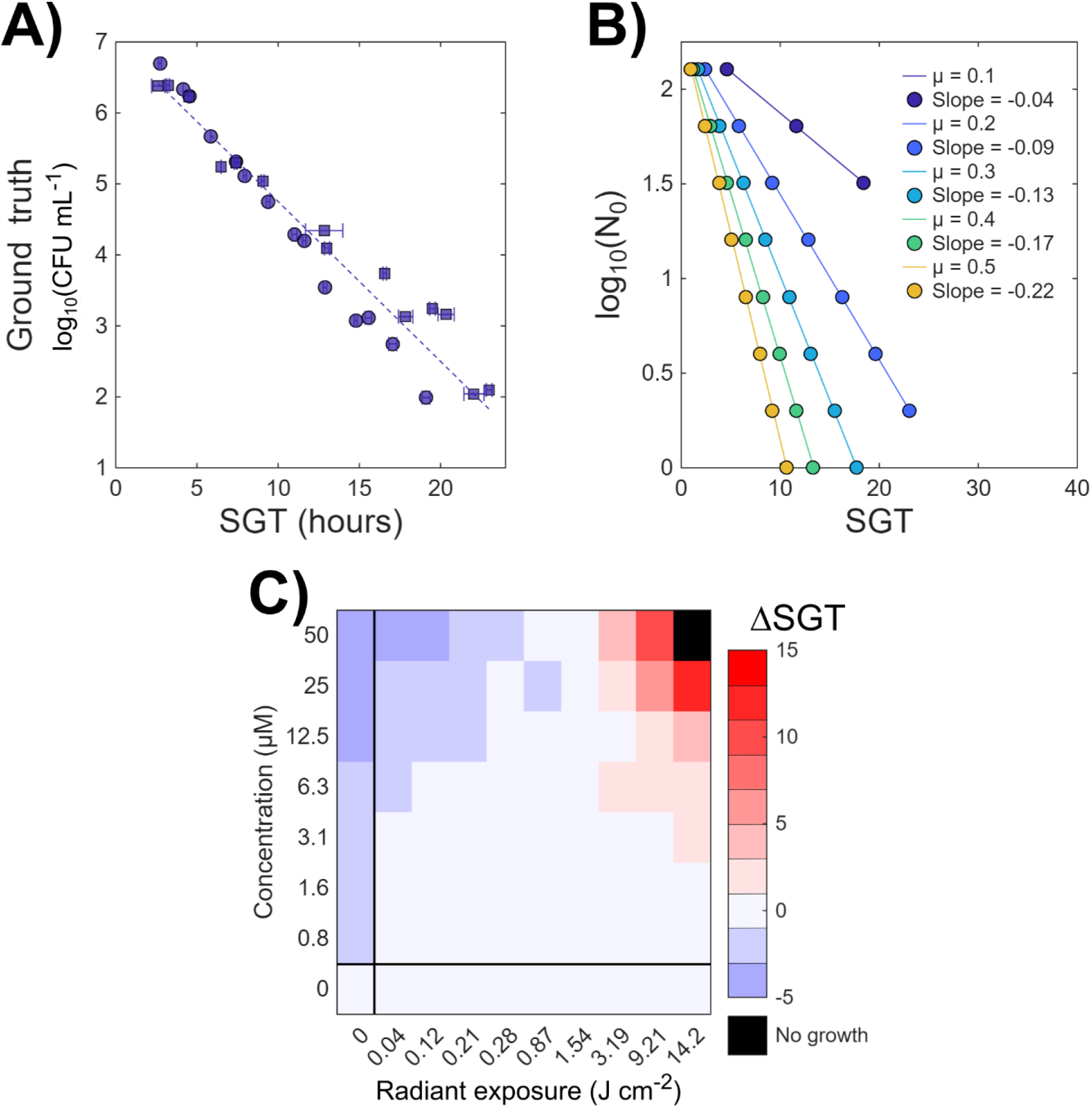
SGT method as a point of comparison. **A)** Starting number of cells as function of the SGT extracted from the dataset used in Figure 2, with SGT defined at *OD*_600_ = 0.05. Markers and error bars depict the mean and standard deviation of three technical replicates, with square and circular markers depicting two independent experiments. **B)** Simulated SGT as a function of decreasing initial number of cells, assuming different growth rates. For further details on this, refer to Supplementary Material 3. **C)** SGT as compared with the controls, expressed as Δ*SGT* = *SGT*_*sample*_− *SGT*_*control*_.

## Conclusions

Our proposed strategy is based on a well-known microbiological concept, where the exponential phase is governed by an exponential equation. However, this can only be applied when we only consider the growth and no chemotherapeutic effect may be observed, which is not the case for most antimicrobial applications. Physical disinfection methods, which include heating, pressure, gamma radiation and in an extensive sense PACT, could be considered as deadly events, where cellular death happens only during the physical treatment. In this sense, the surviving cells may grow back if the conditions are adequate, with the exponential phase being delayed due to the decrease of the starting number of cells. Along this work we have demonstrated how the calculation of the exponential phase predicts the reduction of cells due to a PACT treatment, validating our results with artificially diluted samples. Furthermore, our strategy provides a new parameter to consider for the photodynamic therapy community, which is the growth rate. The growth rate has been greatly overlooked, and the community needs steady evidence that PACT strategies are resistance-proof. In this work, we found that the presence itself of RB has a potential microstatic activity in the dark, which has already been described in the literature (Fiegler-Rudol *et al*. 2025), with this microstatic activity being present also at non-lethal irradiation conditions, as 50 µM RB and 3.19 J cm^−2^ radiant exposure. The acquisition of this knowledge was only possible when applying our strategy as a high-throughput screening checkerboard of both RB concentrations and light exposition. This strategy could be applied for the comparison of efficiency of different photosensitizers, light sources and PS delivery systems, while also holding potential as an alternative quantitative method to assess the efficiency of other physical disinfectant methods.

## Author contributions

N. M.-C.: conceptualization, methodology, software, validation, formal analysis, investigation, data curation, writing – original draft, writing – review& editing, visualization. G. I.: conceptualization, resources, writing – review & editing. G. R.: conceptualization, resources, writing – review & editing, supervision, project administration, funding acquisition. F. F.: resources, supervision, project administration, funding acquisition

## Supplementary data

Supplementary Figures can be found as part of the Supplementary Material, hosted at …

Additionally, an example of the code developed during the current project can be found in the following link: https://github.com/NidiaMC/SupMat1. The provided code is accompanied with the dataset of one experimental day used for the validation of the method. This code can be implemented to any other type of growth curve, adapting the timepoints as convenient.

