## Supplementary Information for "Quantitative evaluation of growth rate and survival of planktonic yeast under dual-parameter photodynamic treatment"

Supplementary Material

Nidia Maldonado-Carmona[a], Giacomo Insero*[a], Franco Fusi[a], Giovanni Romano[a]
[a] Department of Biomedical Experimental and Clinical Sciences “Mario Serio”, Università degli Studi di Firenze, Firenze 50134, Italy
*Corresponding authors: N.M.-C., Department of Biomedical Experimental and Clinical Sciences “Mario Serio”, Università degli Studi di Firenze, Firenze 50134, Italy;. G.I., Department of Biomedical Experimental and Clinical Sciences “Mario Serio”, Università degli Studi di Firenze, Firenze 50134, Italy;.

Keywords: photodynamic antimicrobial chemotherapy;

#### Supplementary Material 1. Example of the validation procedure

An example of the code developed during the current project can be found in the following link: https://github.com/NidiaMC/SupMat1. The provided code is accompanied with the dataset of one experimental day used for the validation of the method. This code can be implemented to any other type of growth curve, adapting the timepoints as convenient.

#### Supplementary Material 2. Supplementary figures

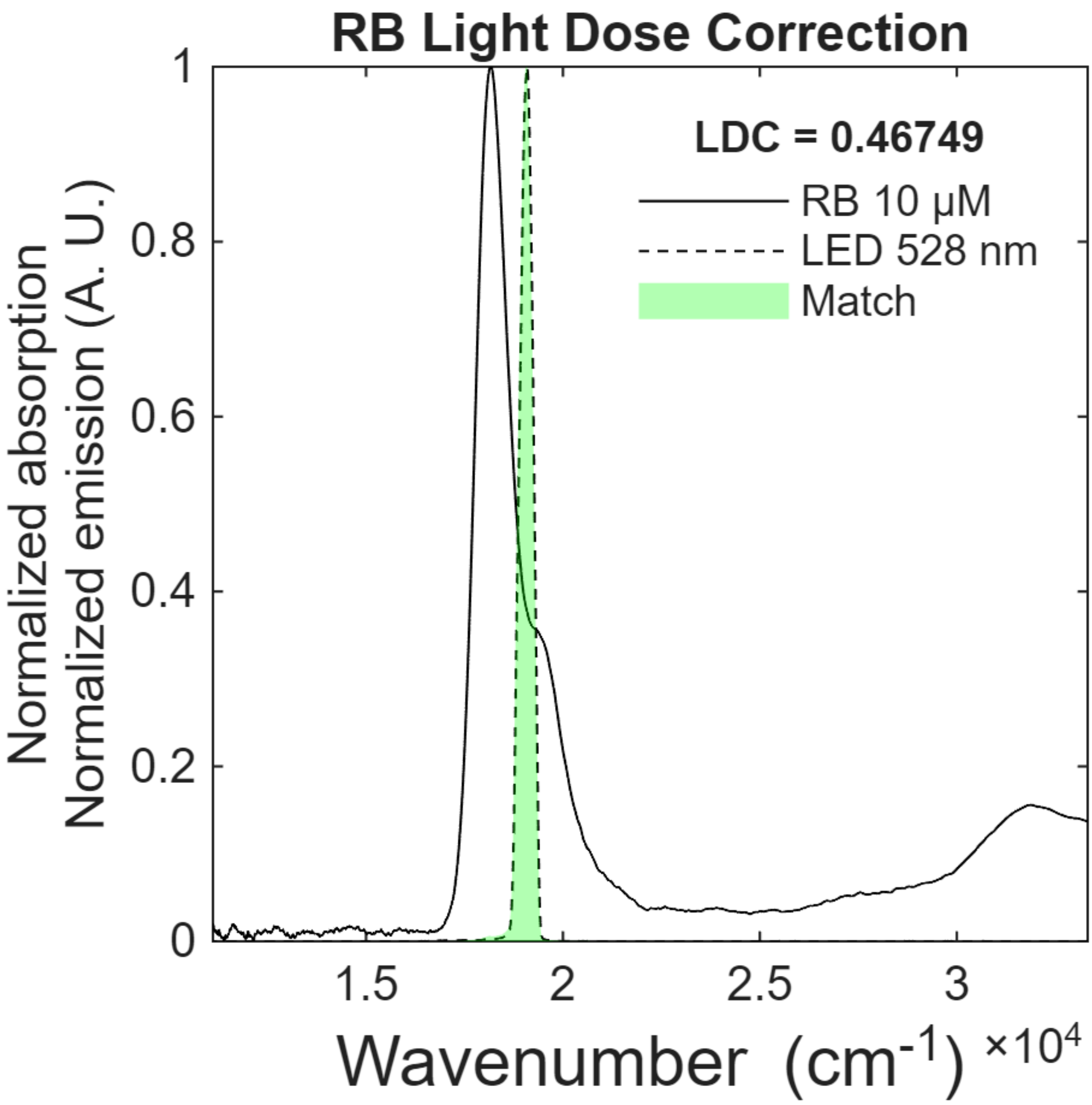


**Supplementary Figure 1**. Match between light source and photosensitizer. Our light source emission spectrum (dashed black line) is compared with the absorption spectra of RB (black continuous line). The match is depicted as the green shadow.

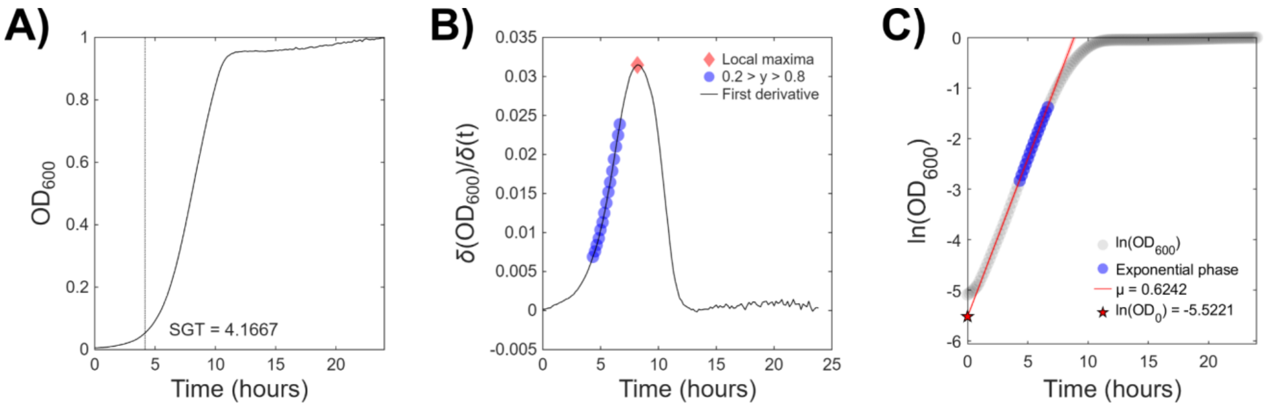


**Supplementary Figure 2**. Extraction of exponential growth parameters from growth curves. **A)** Growth curve expressed as optical density. The SGT where the optical density reaches 0.05 is expressed as a vertical line, which also defines if growth in a sample occurred. **B)** First derivative of the growth curve with respect to time. The maximum growth rate is identified (red diamond) and the exponential phase (blue circles) is considered where the first derivative values are found in between the 20 and 80% of the maximum growth rate. **C)** Semi logarithmic growth rate curve (gray circles) where the exponential phase (blue circles) is fitted to a linear model following the equation $\ln(OD) = \ln(OD_0) + \mu t$. Then, μ is found as the slope (red line), while $\ln(OD_0)$ is found as the intersection with the y-axis (red star).

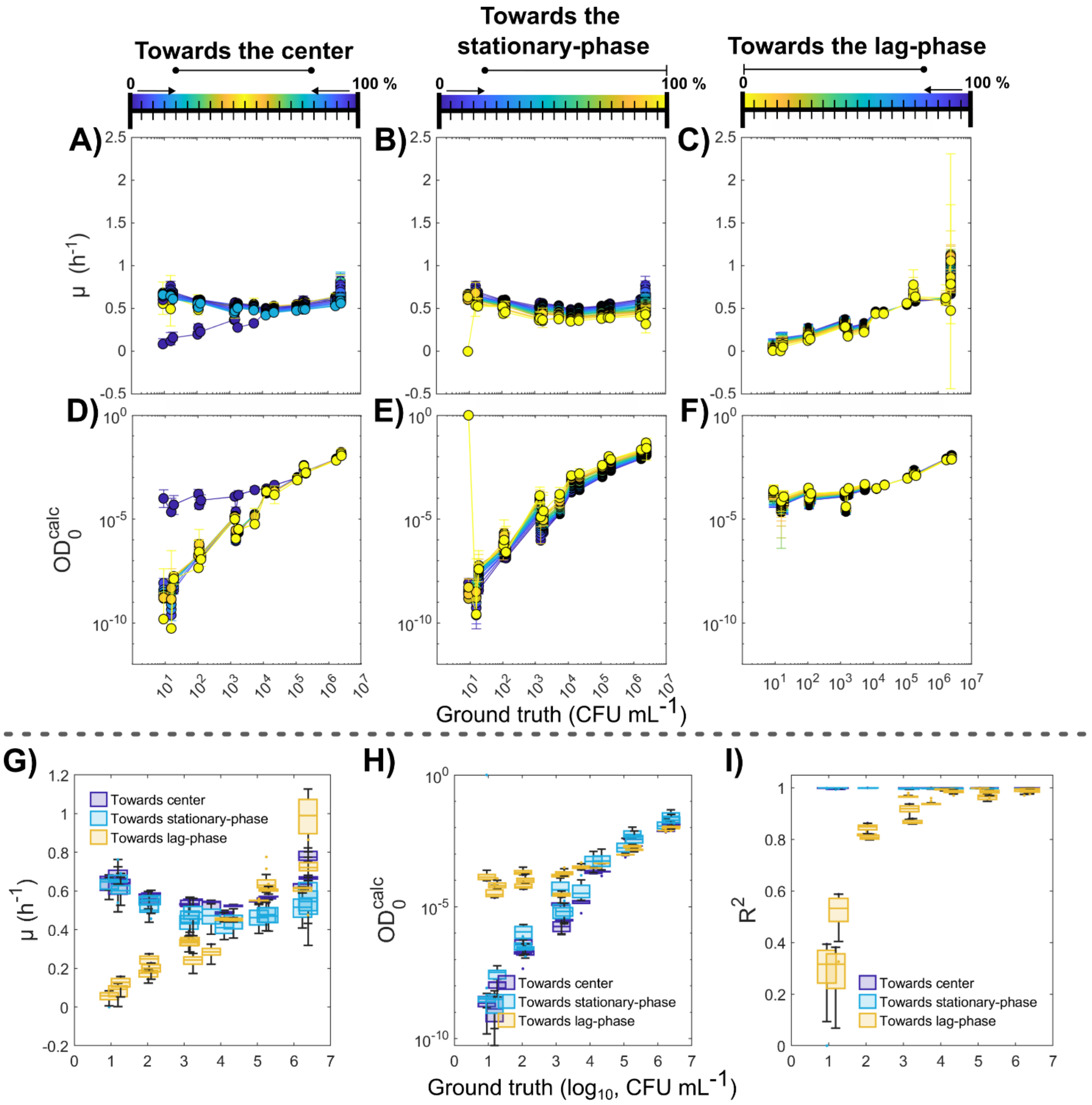


**Supplementary Figure 3**. Determination of growth descriptors with different defined extends for the exponential phase. Growth rate (Panels A, B and C) and $OD_0^{calc}$ (Panels D, E and F) were calculated considering different OD ranges, with respect to the inflexion point found at the δOD / δt curve, as determined in Supplementary Figure 2B. Ranges were considered between 0% (t = 0, beginning of the experiment) and 100% at $t_{max}$, where we observed the inflexion point. The exponential phase was defined considering three scenarios: i) a moving range towards the 50% with respect to the inflexion point, which excludes both the lag and the stationary-phase interfaces (Panels A and D); ii) a moving range towards the stationary-phase (5 to 100%), starting at 5% of the inflexion point magnitude, which excludes the lag-phase interface (Panels B and E); and iii) a moving range towards the lag-phase (0% to 95%), starting at 95% of the inflexion point magnitude, which excludes the stationary-phase interface (Panels C and F). The descriptors were calculated for several starting OD and plotted against their ground truth. To assess the effect of the selection of different extends of the exponential phase, the distribution of the obtained values was plotted as a function of the ground truth (Panels G-I). Although values of μ (Panel G) and $OD_0^{calc}$ (Panel H) group nicely, the predicted values when including the lag-phase interface diverts from the values obtained when ignoring it, with the difference being more striking when decreasing the number of cells, as the extent of the lag-phase increases in time. Additionally, when considering the lag-phase interface as a part of the data to fit, the goodness of fit decreases drastically below 0.9 ($R^2$, Panel I), while ignoring the lag-phase yields $R^2$ values above 0.98 in all cases.

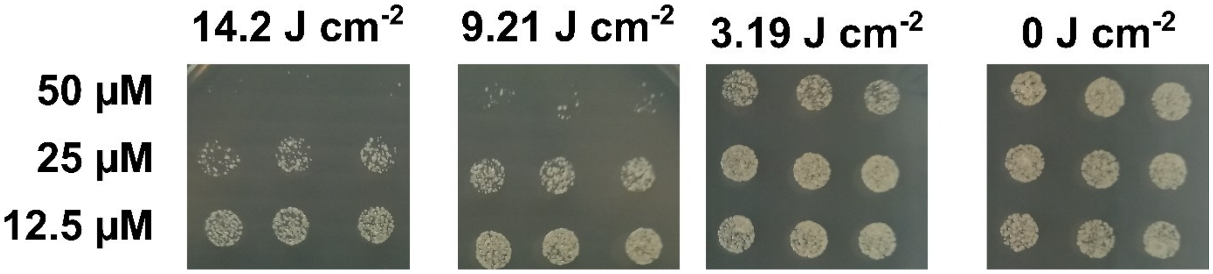


**Supplementary Figure 4**. Decrease of cell viability observed through colony counting spot method. Here we observe that viability quickly decreases as a function of an increase in both light and RB concentration. Every condition is represented by three individual spots, which were plated without previous dilution.

### Supplementary Material 3. Influence of growth rate on SGT determinations

To further explore the relationship between growth rate, initial number of cells and SGT, we simulated the exponential growth of several populations ($N = N_0 e^{\mu t}$), ranging from 0 to 24 hours in 10-minute lapses. To highlight our point, we simulate the growth when starting with different growth rates ($\mu$, from 0.1 to 0.5 $h^{-1}$) and different number of cells ($N_0$, 1 to 128). An example of the growth curves obtained with a fixed number of cells ($N_0$ = 32) and increasing growth rates or with a fixed growth rate ($\mu$ = 0.3) and increasing number of cells can be found in Supplementary Figure 3.1 A and B, respectively. For all the growth curves obtained, the threshold at 200 cells was defined and calculated (Supplementary Figure 3.1 C). We can observe that SGT is greatly affected by the difference on the growth rate of the hypothetical population, observing false negatives for populations with slow growth rates: when the growth rate falls at 0.1, growth is detected in only 38% of the samples (Figure 5B and Supplementary Figure 3.1 C). This indicates that the sensitivity of the method, understood as the capacity to predict the starting number of cells, decreases with decreasing growth rates, while also leading to wrong comparison, when comparing samples with different growth rates but the same starting number of cells.

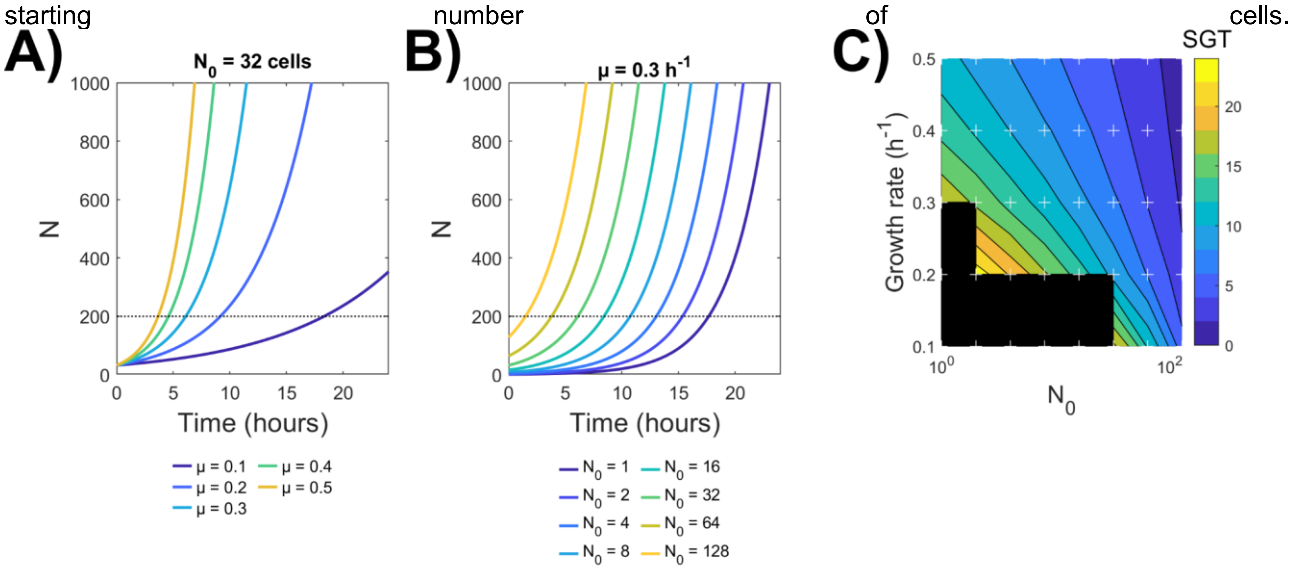


**Supplementary Figure 3.1**. Simulated exponential growth curves, with a fixed starting number of cells ($N_0$ = 32 cells, **A**) or a fixed growing rate ($\mu$ = 0.3 $h^{-1}$, **B**); the pointed line shows the threshold for SGT at $N$ = 200. Curves for all the chosen combinations were calculated, and their found SGT (**C**) was plotted as a function of the starting number of cells and the growth rate. Black boxes correspond to cases where $N$ < 200 at 24h.
